# Implications of gen(om)e duplications on the expansion and evolution of the GPCR signalling pathway

**DOI:** 10.1101/2024.11.25.625254

**Authors:** Ana Barradas, Waldan K. Kwong

**Affiliations:** Faculty of Biology, Medicine and Health, University of Manchester, Michael Smith Building, Oxford Road, M13 9PT Manchester, UK; Gulbenkian Institute for Molecular Medicine, Rua da Quinta Grande, 6, 2780-156 Oeiras, Portugal

**Keywords:** duplication, GPCR, signalling, evolution

## Abstract

Gene and genome duplications are important evolutionary events associated with the emergence of gene families and novel biological functions. G protein-coupled receptors (GPCRs) constitute the largest family of membrane proteins, and their associated signalling pathways control crucial physiological functions such as neurotransmission, endocrine activity, and immunity. However, the duplication history of the entire pathway across evolutionary time is unknown. Here, we performed a comprehensive analysis of the duplication events of the main interactors of the GPCR signalling cascade. We show that different components of the pathway evolved under distinct frequencies of duplication events, with G proteins and GPCRs exhibiting higher frequencies than the downstream mediators and regulators. We also found that GPCRs are evolutionarily younger than G proteins and that most receptors evolved before their ligands. Additionally, the GPCR signalling system experienced significant gene expansion through duplication during the emergence of placental mammals, which played an important role in all human body systems, particularly concerning ligands and G proteins. These results indicate that the expansion and diversification of the GPCR signalling pathway was based on independent and discrete duplication events of its main components, suggesting that the maintenance of duplicate genes within the pathway may have been mediated by the selection of complementary duplication and divergence processes between the signalling components at specific evolutionary stages.

## Introduction

Gene duplication processes are major contributors to the evolution of biological complexity (Lundin, 1999; He and Zhang, 2005). Emerging patterns of shared syntenic loci in completely sequenced genomes have not only strengthened Susumu Ohno’s concept of evolution by gene duplication (Ohno, 1970), but have also evidenced two fundamental scales of duplication instances: mechanisms giving rise to small-scale duplications (SSDs) normally span one or a few genes, or even part of a gene, while large-scale events such as aneuploidy and polyploidy involve the duplication of lengthier portions of genomic DNA and cover entire chromosomes or whole genomes. In both cases, duplication is regarded as having a considerable evolutionary impact on the development of duplicate gene families in eukaryotic organisms (Wolfe and Shields, 1997; McLysaght et al., 2002; Van de Peer et al., 2003; Putnam et al., 2008). G protein-coupled receptor (GPCR) gene families encode the largest superfamily of functionally related membrane proteins and represent a significant class of drug targets implicated in numerous diseases (Overington et al., 2006). In the human genome, they are divided according to the GRAFS classification system into five main families (glutamate, rhodopsin, adhesion, frizzled/taste2 and secretin) that have descended from a common ancestor through tandem, segmental and whole genome duplication (WGD) events (Fredriksson et al., 2003). GPCRs are the main components of a large-scale signalling framework consisting of the dynamic integration of G protein pathways, ancillary subpathways and regulatory cross-communications (Selbie and Hill, 1998; Marinissen and Gutkind, 2001; Neves et al., 2002), and mediate diverse physiological functions including neurotransmission, immunity, and sensory recognition (Sun and Richard, 2012). The canonical GPCR signalling mechanism consists of a linear sequence of information transmission that is initiated upstream of the cell membrane, where extracellular ligands bind to and activate receptors (GPCRs) that intracellularly couple to heterotrimeric guanine nucleotide-binding proteins (G proteins). Activated G proteins, in turn, modulate second messenger-generating systems that further propagate the signal by recruiting secondary effectors that generate biological responses (Figure 1). Notwithstanding, this classical view represents an incomplete and oversimplified depiction of GPCR signalling as together with activating various pathways as the downstream targets of GPCR stimulation such as MAPK and PI3K/Akt cascades, multiple lines of evidence suggest that GPCR signalling plays a vital role in integrating transduction pathways involving other classes of membrane receptors (Rozengurt et al., 2010). The transactivation of receptor tyrosine kinases (RTKs) through GPCR agonists like endothelin and lysophosphatidic acid (Daub et al., 1996; Natarajan and Berk, 2006; Cattaneo et al., 2014) and crosstalk with the serine/threonine kinase receptor family (Burch et al., 2010; Kamato et al., 2015) provide good examples of broader interactions that structure large and complex signalling networks.

**Figure 1.**
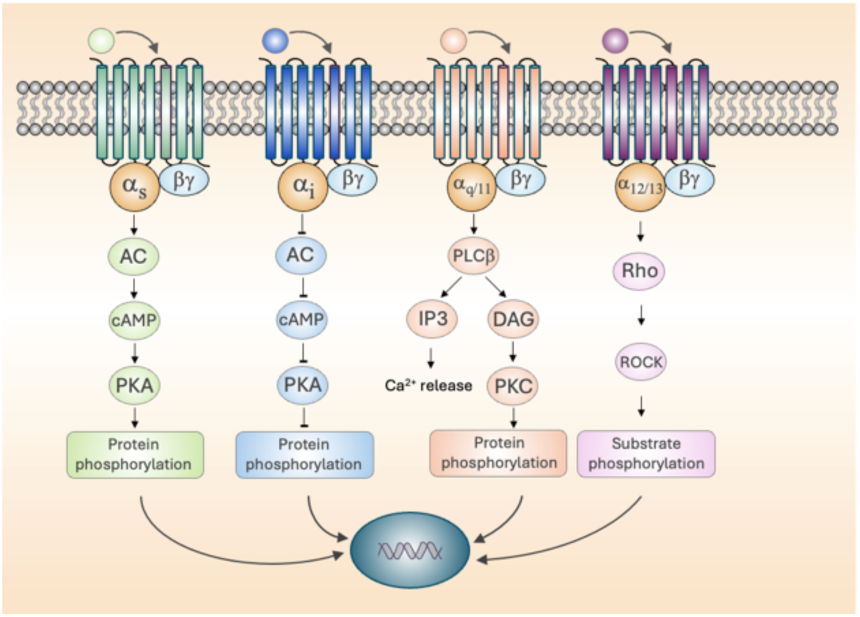
The core GPCR pathway. Downstream signalling pathways of GPCRs based on G protein subunit coupling.

Analyses of protist genomes indicate that protein signalling domains, including a functional GPCR signalling pathway, were already present at the ancestral eukaryote and imply that they played an important role in the origin of metazoan multicellularity (King et al., 2003; Anantharaman et al., 2007; Manning et al., 2008; Suga et al., 2012; de Mendoza et al., 2014). Signalling proteins are also the most representative functional class of duplicates retained after WGD (Huminiecki and Heldin, 2010), reflecting a significant contribution to the evolution of vertebrate complexity (Holland, 1992; Holland, 2003). In contrast to the hypothesised two rounds (’2R’) of WGD (McLysaght et al., 2002; Gu et al., 2002; Dehal and Boore, 2005; Putnam et al., 2008), postulated to have occurred in the stem lineage of vertebrates around 450 million years ago (MYA), SSD is an ongoing and widespread process occurring at genomic and evolutionary scales (Lynch and Conery, 2000; Lynch et al., 2001) and is therefore also likely to affect signalling systems. Duplication processes are widely recognised as underlying the evolution of various GPCR signalling components (Sierra et al., 2002; Indrischek et al., 2017; Mushegian et al., 2012; Alvarez, 2008). Several studies have uncovered important duplication features contributing to the evolution of diverse GPCR subsystems (Braasch et al., 2009; DeVries et al., 2006; Lagerström et al., 2005; Semyonov et al., 2008; Huang et al., 2009; Hwang et al., 2013; Yun et al., 2015; Yamamoto et al., 2012; Tostivint et al., 2014; Gloriam et al., 2005; Cardoso et al., 2010; Valsalan and Manoj, 2014; Yegorov and Good, 2012). However, a global perspective elucidating and quantifying the duplication history of the broader GPCR signalling system has not been carried out to date. In this work, a detailed study of the duplication history of the GPCR signalling system was performed. The analysis revealed that the individual components display different signatures of duplication, suggesting that interactions at specific evolutionary stages were crucial in the growth of the pathway and the evolution of complexity of the signalling system.

## Results

### Identification of signalling components and organisation of the GPCR signalling system

A combination of search methods and manual curation were used to determine and classify the main interactors involved in the human GPCR signalling system. A total of 1,288 proteins implicated in GPCR-mediated signalling were identified, belonging to four distinct signalling components that were broken down into functional subgroups. Each component was defined according to its topological position in the signalling system: activating molecules such as endogenous ligands typically localise extracellularly and upstream of transmembrane GPCRs, whereas the transduction cascade is intracellular and located downstream of G proteins (Figure 1). The overall architecture of the system shows that GPCRs form the largest component, with 372 genes (even without considering olfactory receptors), and that they are activated by a relatively high number of ligands (150 gene-encoded proteins). However, it should be noted that many of the analysed GPCRs are activated by ligands that are not encoded by a gene, (e.g., metabolites, ions, lipids, etc.) and, therefore, Table 1 shows an underestimate of their entire complement of activating ligands. In contrast, GPCRs interact with a relatively small number of G protein isoforms (33 genes), but these activate a substantial amount of downstream effectors (290 genes), thus conforming to the typical bow-tie structure of GPCR signalling (Polouliakh et al. 2009) that consists of numerous inputs and outputs connected through a narrow core, which is also found in other physiological networks (Ma and Zeng 2003; Oda et al. 2005; Oda and Kitano 2006).

**Table 1.** Overview of the duplication history features of the human GPCR signalling system components.

| Topological position | Signalling component | Number of genes | Number of duplicates / singletons | Duplication events | Most frequent duplication (MYA) | EDN (MYA) | LDN (MYA) | ESN (MYA) |
| --- | --- | --- | --- | --- | --- | --- | --- | --- |
| GPCRs | GPCRs | 372 | 352 / 20 | 1906 | Euteleostomi (441) | Bilateria (937) | *Homo sapiens* (0) | Bilateria (937) |
|  | Adhesion | 32 | 32 / 0 | 189 | Chordata (722) | Bilateria (937) | Hominoidea (20) | - |
|  | Frizzled | 11 | 11 / 0 | 69 | Bilateria (937) | Bilateria (937) | Eutheria (104) | - |
|  | Secretin | 15 | 15 / 0 | 102 | Bilateria (937) | Bilateria (937) | *Homo sapiens* (0) | - |
|  | Rhodopsin | 289 | 273 / 16 | 1464 | Euteleostomi (441) | Bilateria (937) | *Homo sapiens* (0) | Bilateria (937) |
|  | Glutamate | 19 | 18 / 1 | 78 | Bilateria (937) | Bilateria (937) | Euarchontoglires (92) | Chordata (722) |
|  | 7TM orphan | 6 | 3 / 3 | 4 | Euteleostomi (441) | Euteleostomi (441) | Euteleostomi (441) | Bilateria (937) |
| Upstream | Ligands | 150 | 82 / 68 | 321 | Euteleostomi (441) | Bilateria (937) | *Homo sapiens* (0) | Vertebrata (535) |
|  | Rhodopsin ligands | 113 | 59 / 54 | 222 | Euteleostomi (441) | Bilateria (937) | *Homo sapiens* (0) | Euteleostomi (441) |
|  | Secretin ligands | 18 | 4 / 14 | 12 | Eutheria (104) | Euteleostomi (441) | Eutheria (104) | Vertebrata (535) |
|  | Frizzled ligands | 19 | 19 / 0 | 87 | Bilateria (937) | Bilateria (937) | Euteleostomi (441) | - |
|  | Chemokine | 39 | 27 / 12 | 113 | Theria (162) | Euteleostomi (441) | *Homo sapiens* (0) | Euteleostomi (441) |
|  | Complement component | 2 | 2 / 0 | 14 | Chordata (722) | Bilateria (937) | Amniota (296) | - |
|  | Hormone | 8 | 7 / 1 | 32 | Euteleostomi (441) | Euteleostomi (441) | *Homo sapiens* (0) | Euteleostomi (441) |
|  | Neuropeptide | 56 | 13 / 43 | 37 | Euteleostomi (441) | Euteleostomi (441) | Catarrhini (29) | Euteleostomi (441) |
|  | Peptide/protein | 45 | 33 / 12 | 125 | Bilateria (937) | Bilateria (937) | *Homo sapiens* (0) | Vertebrata (535) |
| Downstream | G proteins | 33 | 32 / 1 | 208 | Bilateria (937) | Bilateria (937) | Catarrhini (29) | Vertebrata (535) |
| | Gα | 16 | 16 / 0 | 126 | Bilateria (937) | Bilateria (937) | Eutheria (104) | - |
| | Gβ | 5 | 5 / 0 | 31 | Bilateria (937) | Bilateria (937) | Eutheria (104) | - |
| | Gγ | 12 | 11 / 1 | 51 | Euteleostomi (441) | Euteleostomi (441) | Catarrhini (29) | Vertebrata (535) |
| | G$\alpha_s$ | 2 | 2 / 0 | 26 | Bilateria (937) | Bilateria (937) | Eutheria (104) | - |
| | G$\alpha_i$ | 8 | 8 / 0 | 62 | Bilateria (937) | Bilateria (937) | Euteleostomi (441) | - |
| | G$\alpha_{q/11}$ | 4 | 4 / 0 | 24 | Bilateria (937) | Bilateria (937) | Euteleostomi (441) | - |
| | G$\alpha_{12/13}$ | 2 | 2 / 0 | 14 | Bilateria (937) | Bilateria (937) | Amniota (296) | - |
|  | Transduction cascade | 290 | 289 / 1 | 1570 | Bilateria (937) | Bilateria (937) | *Homo sapiens* (0) | Euteleostomi (441) |
| Peripheral | Regulation | 145 | 127 / 18 | 624 | Bilateria (937) | Bilateria (937) | Catarrhini (29) | Opisthokonta (1215) |
|  | Crosstalk | 298 | 243 / 55 | 1236 | Bilateria (937) | Bilateria (937) | *Homo sapiens* (0) | Opisthokonta (1215) |
| Protein-coding genes |  | 19784 | 14271 / 5513 | 68519 | Bilateria (937) | Opisthokonta (1215) | *Homo sapiens* (0) | Opisthokonta (1215) |

EDN – Earliest duplication node; LDN – Latest duplication node; ESN – Earliest speciation node; MYA – Million Years Ago; Phylogenetic dating follows Ensembl (version 84) timing.

### The components of the GPCR pathway show different duplication patterns

A comprehensive analysis of the extent and pattern of duplication of the topological layers of the GPCR signalling system was performed by comparing the duplication and speciation frequencies of each signalling component with those of all protein-coding genes at each evolutionary clade (Figure 2). Three evolutionary stages can be distinguished comprising an early time period between the emergence of bilaterians and bony vertebrates (937-441 MYA), an intermediate phase around mammalian radiation (371-104 MYA), and a recent stage between the anthropoid ancestor and human speciation (42-0 MYA). The duplication/speciation events of the signalling components distribute non-randomly across and within these stages, producing distinct duplication signatures. G proteins expanded the most around the emergence of bilaterians, while GPCRs and the intracellular signalling machinery experienced continued duplication events from the emergence of bilaterians until the time of WGD (Euteleostomi). Gene-encoded ligands, in turn, were duplicated more frequently at the intermediate and recent stages. Moreover, tracing the duplication origins of the signalling components as their earliest duplication node (EDN) supports a pattern of evolutionary transitions in the development of G protein signalling, suggesting that the ancient intracellular origins of this function did not require membrane receptors. Our results suggest that the system was subsequently expanded by gene and genome duplications that diversified GPCRs and at later stages, their activating ligands (Figure S1C).

**Figure 2.**
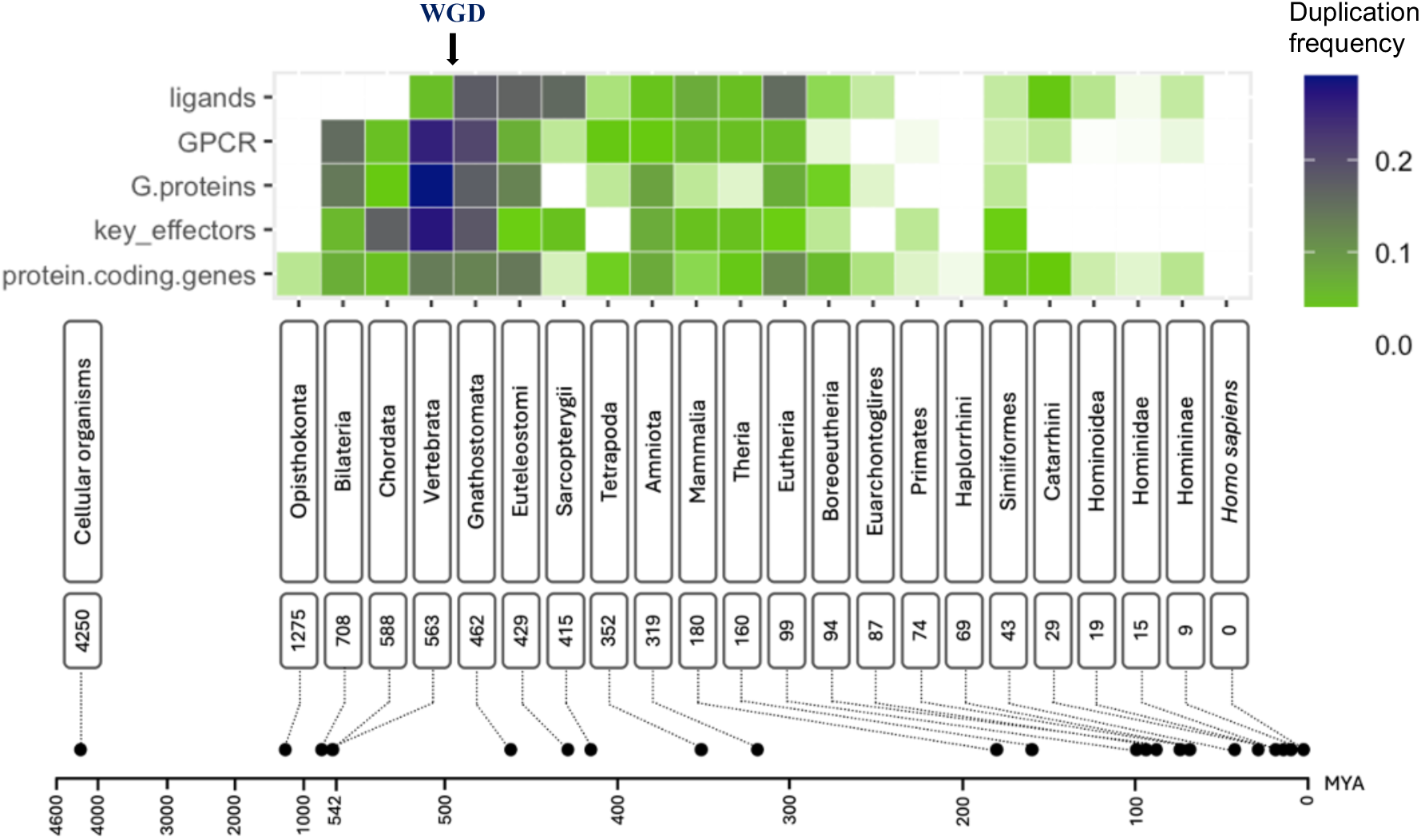
Duplication history of the components of the GPCR signalling system. The heatmap represents duplication ratios for each clade across evolutionary time. **WGD** - Whole genome duplication.

In order to assess the evolutionary trajectory of duplicated genes within the GPCR signalling system, we assigned to each duplicate gene its latest duplication node (LDN). We observe two clusters of retained genes at the time of WGD (Euteleostomi) and Eutheria, indicating that signalling by GPCRs played an important role in the development of vertebrates and placental mammals (Figure S1B). We have also traced the origin of singleton genes within the pathway as their earliest speciation node (ESN) through their speciation events. We found that singleton origins tend to be more recent than duplication ancestries and also that unduplicated genes tend to be found within the ligand component (Figure S1D). These observations indicate that fine-tuning of GPCR signalling by activating ligands is a relatively recent innovation that was achieved by exploitation of both duplicate and singleton genes, especially within chemokines and neuropeptides.

We also evaluated whether individual GPCR families and their cognate ligands follow the same trend in their duplication history. Interestingly, we found that GPCR GRAFS families were expanded sequentially, with frizzled and secretin families reaching the highest peaks of duplication at the emergence of bilaterians, adhesion expanding at the origin of chordates, rhodopsin duplicating most frequently at the base of vertebrates, and glutamate peaking at the expansion of bony vertebrates (Figure S1A). Similarly, there is also a sequential, but more recent, duplication trend in cognate ligands. Frizzled ligands (Wnt proteins) are the most ancient and resilient to small-scale duplication, peaking at Bilateria and confirming the importance of the Wnt pathway in the establishment of polarity during metazoan embryonic development (Petersen and Reddien 2009). Secretin ligands were subsequently expanded by WGD and SSD at the emergence of placental mammals, and followed by rhodopsin ligands, which were expanded by duplication more recently through SSD at the intermediate and recent stages. We also detected a slight consecutive pattern of duplication within G protein subunits, with duplication common ancestries at Bilateria for *alpha* and *beta* subunits, and Euteleostomi for *gamma* subunits (Figure S1A). *Alpha* subunits show the most ancient LDNs, with the exception of G_αs_, followed by more recent LDNs of *beta* and *gamma* subunits (Figure S1B).

Taken together, these observations indicate that gene/genome duplication was a major mechanism of evolution of the GPCR signalling pathway and that the different components evolved under distinct duplication rates that were not constant over evolutionary time. Furthermore, expansion of the system was also achieved by single copy ligands and proteins that promote crosstalk with other pathways.

### Duplication history and interacting partners

When compared to GPCRs that are activated by gene-encoded ligands, GPCRs that are activated by metabolites have preferentially been retained after WGD in the pufferfish genome (Semyonov et al., 2008). In contrast, human GPCRs whose cognate ligands are gene-encoded, have been shown to have higher duplication frequencies in more recent evolutionary stages than those that are activated by non-peptide ligands (Huang et al. 2009). Having established significant differences in evolution by duplication between the main components of the signalling system, we hypothesized that the duplication history of the receptors might correlate with their diversification in terms of functional features. Therefore, we investigated the relationships between interacting pairs of GPCRs and cognate ligands, and GPCRs and coupling G proteins.

Firstly, GPCRs were grouped into four categories, depending on their type of activating ligand (metabolite, gene-encoded, sensory or orphan) and their evolutionary properties were compared in terms of duplication rate and tissue expression (Figure 3). There was no evidence to support an overall association between the number of ligands (or receptors) a GPCR (or ligand) binds and their underlying duplication history or tissue specificity (Table 2). No differences were found regarding tissue specificity as assessed by the index τ, however, there is a significant difference (*P* = 0.002, Kruskal-Wallis test) across all four groups when considering simple tissue counts (Table 2). Dunn’s test (*P* < 0.001) further showed that the sensory GPCRs are the most specific receptors regarding the number of tissues where they are expressed, which is compatible with photoreceptor gene expression in rod and cone cells. Regarding the number of duplication events, there is also a statistically significant difference between GPCRs that are activated by different kinds of ligands, revealing that sensory receptors have experienced higher frequencies of duplication retention across evolution than orphan receptors (*P* = 0.009, post hoc Dunn’s test), which may provide clues to their deorphanization. While there are no differences regarding the duplication frequencies of metabolite-activated GPCRs, some subgroups seem to have diverged considerably regarding their expression patterns. This is the case of aminergic GPCRs and receptors activated by amino acids, which are very tissue-specific, as opposed to the broadly expressed receptors activated by lipids and nucleotides (Figure S2). Differences regarding tissue-specificity were also found within GPCRs activated by gene-encoded ligands, especially between chemokine and frizzled (most broadly expressed), and neuropeptide receptors, which are largely confined to central and peripheral nervous system tissues (Figure S3).

**Figure 3.**
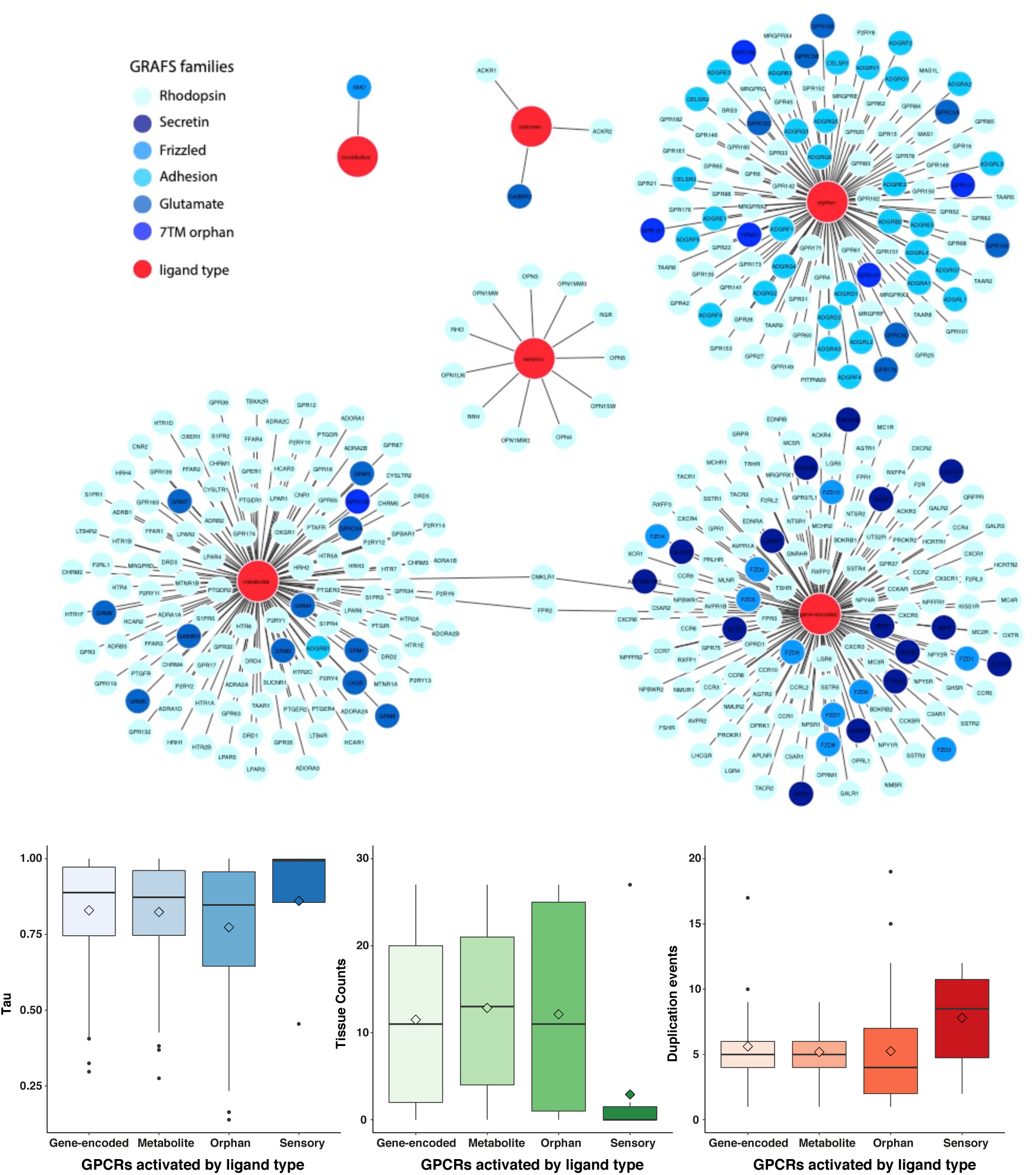
(A) Tissue specificity and duplication events for GPCR-ligand binding interactions. Edges represent binding interactions between ligands and their cognate GPCRs. GPCR nodes are coloured according to GRAFS families and ligand specificities are coloured red. **(B)** Boxplots show tissue specificity: (**1)** τ; (**2)** tissue counts; and (**3**) duplication events for GPCRs with different ligand binding properties. Solid lines inside the boxes indicate the median and diamonds indicate the mean.

**Table 2.**
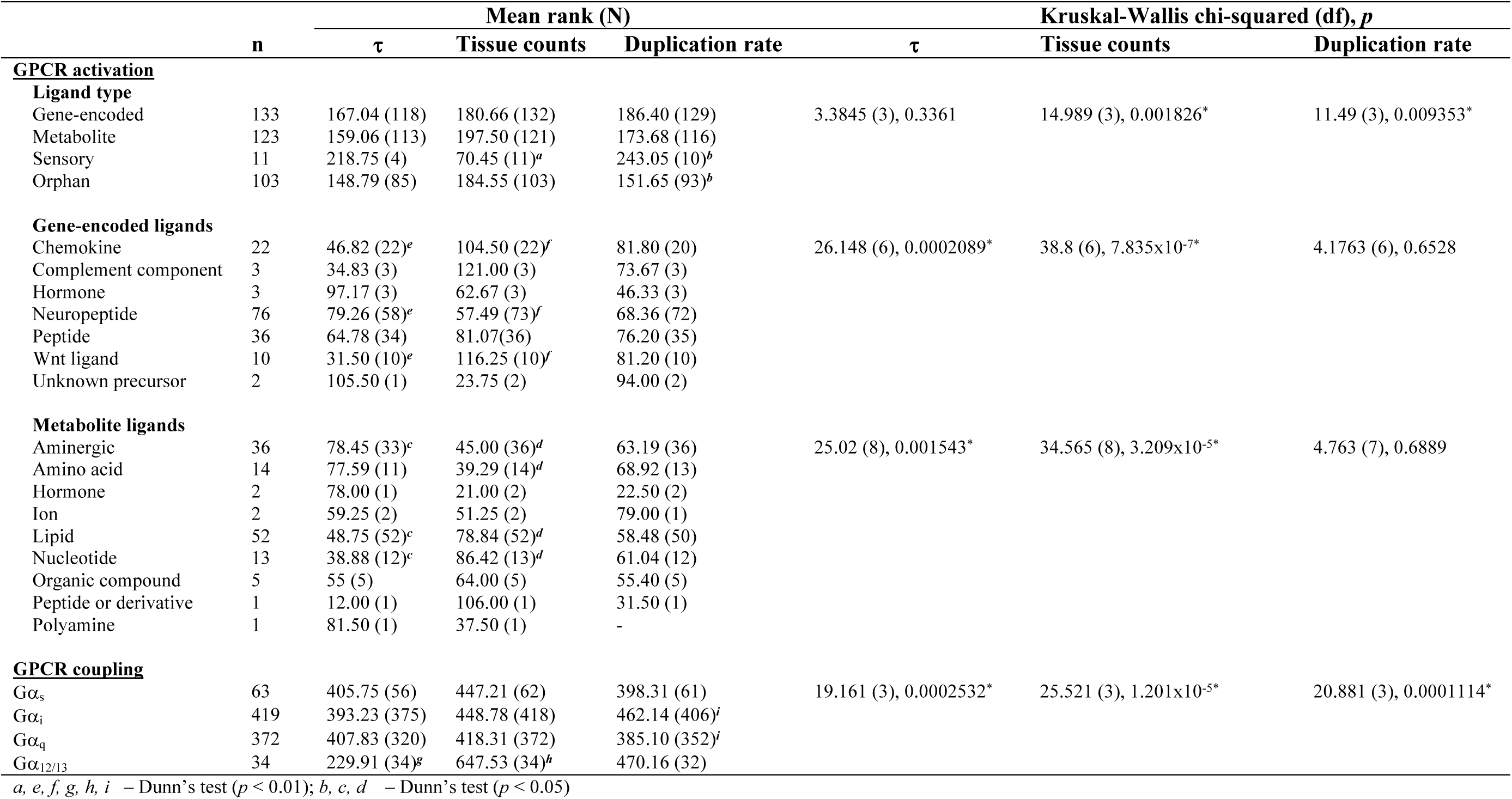
Overview of the differences between GPCR groups in relation to tissue specificity (τ and tissue counts) and duplication rate. Stars indicate significant differences for e Kruskal-Wallis test and significant post hoc comparisons with Dunn’s test are indicated with superscripts.

Secondly, GPCRs were evaluated in terms of their G protein coupling preferences and grouped into four categories; Gα12/13, Gαi, Gαq/11, and Gαs (Figure 4).

**Figure 4.**
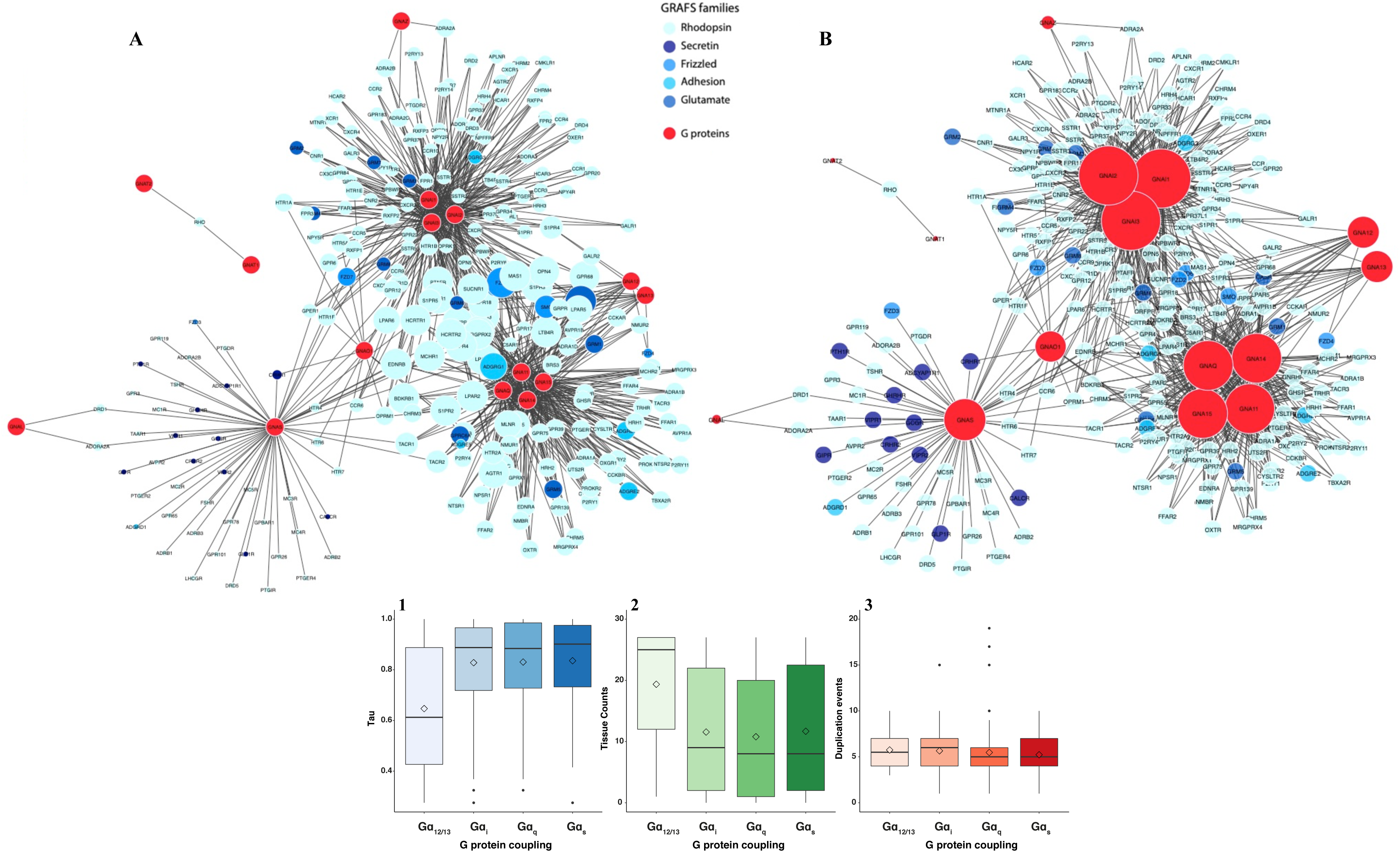
GPCR-G protein coupling specificities. Edges represent receptor- Gα protein coupling pairs in primary transduction mechanisms. GPCR nodes are coloured according to GRAFS families and G protein α subunits are coloured red. **(A)** GPCR node size is proportional to the number of G proteins coupling to the receptor. **(B)** G protein node size is proportional to the number of receptors coupling to each Gα subunit. **(C)** Boxplots show tissue specificity: (**1)** τ; (**2)** tissue counts; and (**3**) duplication events for each Gα protein-coupling GPCR group. Solid lines inside the boxes indicate the median and diamonds indicate the mean.

Based on the correlation analysis between the node degrees of each GPCR coupling preference and tissue specificity and duplication rate we also could not find an association between a G protein coupling preference and their underlying duplication history or tissue specificity (Table 3). However, the more receptors a G protein is able to couple, the more broadly it will be expressed (Spearman’s rank correlation coefficient, r= 0.7422, *P* = 0.0015). Concerning specificity in tissue expression, we observe a contrast depending on the primary transduction mechanisms of the receptors; GPCRs that primarily couple to G_α12/13_ are much more broadly expressed than any other G protein *alpha* subunit coupling (Dunn’s test, P < 0.01), but in spite of this difference, the number of duplication events for G_α12/13_- coupling GPCRs was not significantly higher. However, G_αi_-coupling GPCRs have duplicated significantly more than GPCRs whose transduction mechanism is G_αq/11_ (Dunn’s test, P < 0.01). GPCRs that primarily couple to G_α12/13_ are involved in regulating cytoskeletal remodeling, cell migration, and cellular proliferation, which are required functions in almost every cell type (Suzuki et al., 2009). Receptors that activate the G_αi_ subunit inhibit adenylate cyclase and are typically involved in inhibitory functions, such as reducing neurotransmitter release or heart rate (Watts and Neve, 2005), whereas G_αq/11_-coupling GPCRs activate phospholipase C beta and are more involved in stimulatory functions, such as smooth muscular contraction and glandular secretion (Harden et al., 2011). This may indicate that duplication contributed to the diversification of GPCR functions by allowing functional inhibition. GPCRs coupling to G_αs_ are more tissue-specific and although their duplication rate was not higher, the duplication of this G protein subunit occurred at later evolutionary stages (Figure S1B).

**Table 3.** Correlations between node degrees and tissue specificity and duplication history in subnetworks of ligand binding and G protein coupling.

| Node degree | Variable | $\rho$ | $P$ |
| --- | --- | --- | --- |
| GPCR (ligand) | $\tau$ | -0.0709 | 0.4457 |
|  | Tissue counts | 0.0911 | 0.2968 |
|  | Duplication rate | 0.1258 | 0.1539 |
| Ligand | $\tau$ | -0.0074 | 0.9379 |
|  | Tissue counts | 0.0378 | 0.662 |
|  | Duplication rate | 0.0041 | 0.9743 |
| GPCR (G protein) | $\tau$ | -0.0343 | 0.6133 |
|  | Tissue counts | -0.0055 | 0.9319 |
|  | Duplication rate | 0.0295 | 0.6501 |
| G protein | $\tau$ | -0.6456 | 0.0093* |
|  | Tissue counts | 0.7422 | 0.0015* |
|  | Duplication rate | 0.0459 | 0.8709 |

\* $P < 0.05$

### Evolutionary origin of tissue gene expression

We independently evaluated the evolutionary origin of gene expression in each component of the GPCR signalling pathway across the 11 anatomical MeSH descriptors. Our analysis indicates that there are important differences in the evolutionary ancestry of baseline gene expression patterns within each signalling component (Figure 4). When considering duplicates, GPCR genes have evolutionary origins coinciding with their latest duplication node (LDN) at the base of vertebrates and bony vertebrates in all body systems, indicating their importance in vertebrate innovation, whereas ligand and G protein genes are of eutherian origin, especially in the cardiovascular, hemic and immune, nervous, and stomatognathic systems (Figure 5A). The transduction cascade, regulation and crosstalk components also share an early ancestry for their LDN across all human tissues, however, there are more genes in these components with evolutionary origins at the intermediate (Sarcopterygii - Haplorrhini) and recent (Simiiformes – *H. sapiens*) stages than GPCRs (Figure 5A). Of note, transduction cascade genes originating at the last therian ancestor have an important genetic expression across all tissues which is not seen in any other component. For genes involved in crosstalk with other pathways, the two rounds of WGD around the Euteleostomi clade were particularly important for the cardiovascular system, whereas the most recent LDN of *Homo sapiens* represents an important contribution to the total gene expression of embryonic structures (Figure 5A). The regulatory genes have bony vertebrate, amniote and mammalian origins across all body systems, and this is the component with the highest expression of genes of catarrhinian ancestry (Figure 5A).

**Figure 5.**
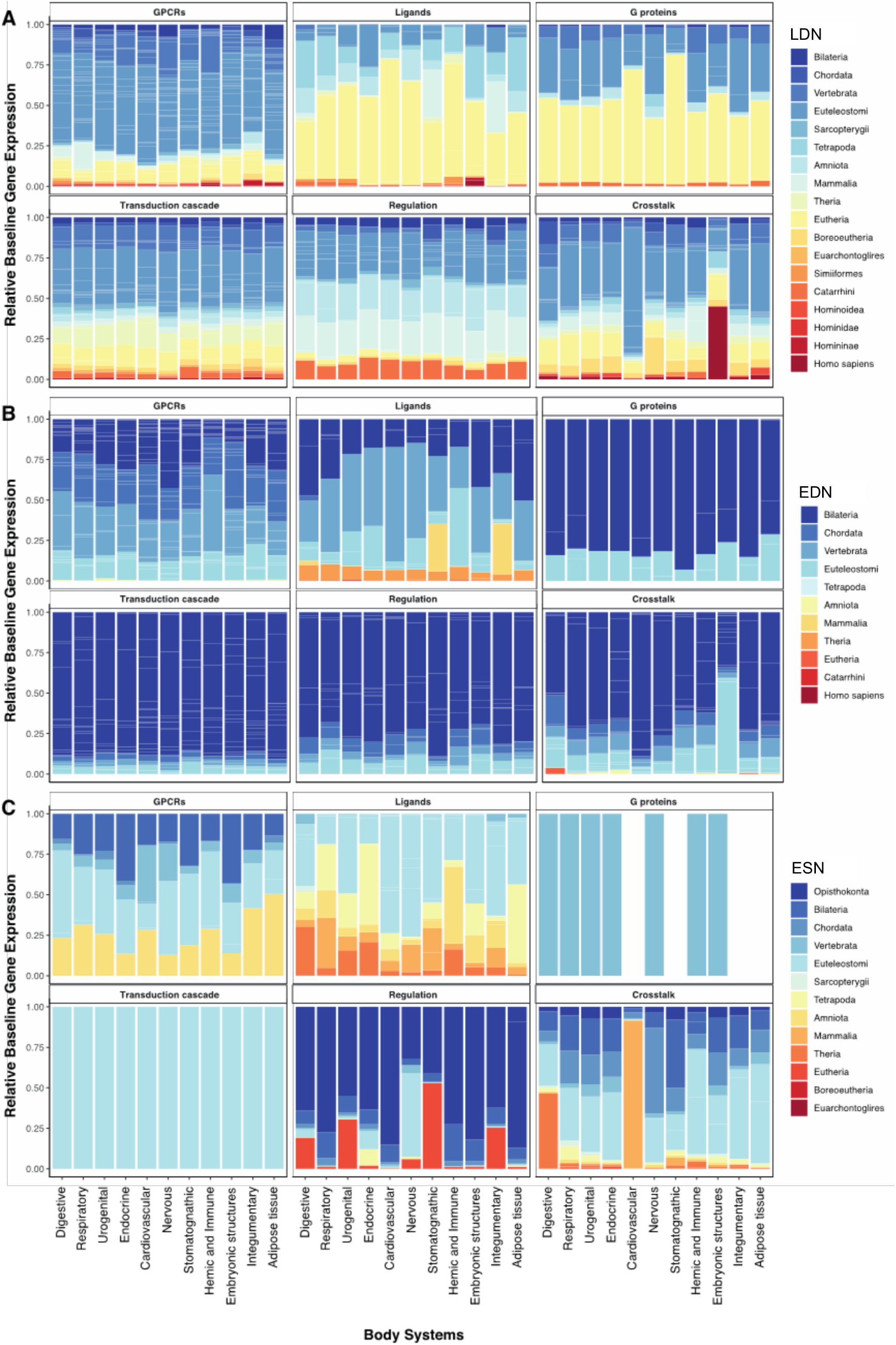
Evolutionary origins of gene tissue expression within the components of the GPCR signalling system. **(A)** Latest duplication node (LDN), **(B)** Earliest duplication node (EDN), and **(C)** Earliest speciation node (ESN) of relative baseline gene expression of genes in the human body systems.

When considering the earliest ancestor of each duplicate gene (EDN) we find that, in most signalling components, gene expression is primarily composed of evolutionarily ancient genes dating from the early emergence of bilaterians, with the exception of GPCRs, whose evolutionary origins are of Bilateria-Euteleostomi descent, and ligands which have the most recent and divergent ancestries extending from Bilateria to Theria (Figure 5B).

We also considered the gene expression in the different body systems of the earliest ancestor of singleton genes (ESN) in the signalling system and identified not only very different patterns of singleton evolutionary origin within the components of the pathway, but also more inconsistent signatures within the different tissues when compared to their duplicate counterparts (Figure 5C). Singleton GPCRs are in their majority Euteleostomi-old, possibly reflecting genes that have lost their duplicate pairs, but also of bilaterian and amniote origin, while the ligand component shows the highest proportion of genes dating from the intermediate clades spanning Euteleostomi and Theria. The regulatory component has the highest fraction of ancient genes originating from the ancestral opisthokont, with some important contributions from the more recent boreoeutherian ancestor, indicating that the co-option of singleton genes that contributed to the complexity of the system was primarily ancient. Most singleton crosstalk genes date from the euteleostomian clade in all human tissues, with a few exceptions in the digestive, cardiovascular and nervous systems.

## Discussion

Evolution by duplication and divergence allows for functional innovation (Nguyen et al., 2014; Mattenberger et al., 2017). Here, we investigate the duplication history of the GPCR signalling system, which has played a key role in the evolutionary success of GPCRs, making them one of the most diverse protein families in eukaryotes. GPCRs are the largest component of the canonical signalling system, which exhibits the typical arrangement of an hourglass structure (Csete and Doyle, 2004) with different duplication patterns for the distinct signalling units, indicating that different bursts of expansion throughout evolutionary time have gradually scaled increasing inputs and outputs while maintaining a narrow core set of proteins.

The timings of the duplication events that underlie the pathway’s widespread functional diversification were not constant throughout evolution and were most abundant at three discrete time periods comprising early, intermediate and recent evolutionary stages in the human phylogeny (Figure 2); namely between Bilateria and Euteleostomi (early), Eutheria (intermediate), and to a lesser extent, in the time period between Simiiformes and Homininae clades (recent). The highest bursts of duplication seem to have occurred at the early stage, between the origin of bilaterians and WGD in the early vertebrate lineage, and were followed by progressively lower small-scale duplication frequencies up until human speciation. This trend is also observed for the whole set of protein-coding genes in the human genome (Figure S1A), where periods of intense duplication can be seen at the bilaterian ancestor, early vertebrates, placental mammals, and anthropoid primates. These results are consistent with previous studies detailing evolutionary origins of human genes (Domazet-Lošo and Tautz, 2008; Surcel et al., 2008). However, a more detailed analysis of genome evolution (Lewis and Dunn, 2018) suggests that the apparent duplication burst observed at the Bilateria node may actually be artefactual and result from the clumping of deeper phylogenetic branches.

The Ensembl Compara database is predominantly focused on vertebrates (Vilella et al., 2009) including only three non-vertebrate species: fly, worm and yeast. This biased sampling towards vertebrates and a less representative sampling at the root of the tree is likely to affect the estimation of duplications at the base of the tree, for example by inflating the number of duplications at the Bilateria node. Similar issues have recently been discussed (Lewis and Dunn, 2018) regarding gene family gains and losses along the branch of the evolutionary tree giving rise to animals in two studies with significantly different taxonomic samplings: Paps and Holland (2018) found considerable expansion of gene families along the Metazoa stem, while Richter et al. (2018) found a burst of gene family expansion in Choanoflagellata rather than in Metazoa, due to the higher number of choanoflagellate species analysed in their study. Because the Ensembl Compara Gene Trees have a biased taxon sampling, this may be problematic for the resolution of the relationships at the root of the species tree. In particular, the leap from opisthokonts directly to protostomes and deuterostomes may affect the inference of early duplications, especially because a substantial amount of GPCR diversity arose before animals, but after the yeast-animal split (de Mendoza et al., 2014).

We also identified differences in the duplication patterns of the signalling components. GPCR duplications mainly occurred in the early evolutionary stages, whereas ligand duplications, except for frizzled ligands, are relatively younger, showing more intense duplication at the intermediate stage and a more recent wave of duplication (Figure S1A). In contrast, downstream components, especially G proteins, are predominantly ancient and practically lack recent duplications. The order of these duplication events and the dating of the singleton ligand genes suggests a shift in the canonical pathway regulation from bottom-up to top-down, in which the ligands became the agents that controlled the activation of the pathway. Another fact supporting this hypothesis is that the GPCR internal signalling machinery is ancient (de Mendoza et al., 2014) and devoid of singletons (Figure S1A and C).

A slight transition in the duplication peaks within GPCR GRAFS families was also observed (Figure S1A), suggesting a stepwise evolution of the different GPCR families. Our results indicate that the frizzled and secretin receptors are the GPCR families with the most ancient duplication nodes, followed by the adhesion and glutamate receptors, and finally the rhodopsin family. Frizzled receptors are very important in embryonic development and cell polarity, but also play important roles in adult tissues (Huang and Klein, 2004). Evidence suggests that the last common ancestor of cnidarians and bilaterians might already possess four frizzled receptor genes (Schenkelaars et al., 2015), which have later duplicated and diversified into the current set of frizzled receptors. This study suggests that this is the most ancient GPCR family to diversify by duplication and, most remarkably, the only one with ancient duplicate ligands. Secretin ligands, on the other hand, are predominantly singletons, and their eutherian origins indicate that they were especially important to the development of placental mammals. In fact, the peptidergic system plays a crucial role in the development and function of the placenta, primarily by regulating key processes like angiogenesis (Malassiné et al., 1992, Wang et al., 2022), trophoblast invasion (Dumolt et al., 2023), immune tolerance (Paparini et al., 2019), and nutrient transport (Lager and Powell, 2012).

Adhesion GPCRs show a significant increase in duplication at the chordate clade, where there are high levels of duplication ancestry (Figure S1C) and latest duplication nodes (Figure S1B), which is not seen in the other GRAFS families. There is indeed indication that members of the family of adhesion GPCRs were already present in early-branching Chordata such as the lamprey (*Petromyzon marinus*), the lancelet (*Branchiostoma belcheri*) and the sea squirt (*Ciona intestinalis*) (Wittlake et al., 2021), and our study suggests that these receptors are small-scale duplicate genes. Adhesion receptors are particularly important in cell-cell and cell-matrix mechano-sensing (Wilde et al., 2022) and have important roles within the immune system (Tseng et al., 2023). Their emergence within chordates may have played a key role in cellular communication and tissue organization that has been conserved throughout evolutionary history.

Glutamate receptors show a significantly increased number of duplications at the basis of vertebrates, although their duplication ancestry and LDNs are not particularly enriched for this clade (Figure S1A and B). Glutamate receptors are crucial components of the nervous system in vertebrates, serving several important functions that are essential for neuronal communication and plasticity (Niswender and Conn, 2010). Our results indicate that the duplication of glutamate receptors within the Vertebrata clade was fundamental to the evolution of the nonmammalian vertebrate nervous system (Budd, 2015), thereby enabling increased behavioral complexity.

Rhodopsin GPCRs, the most numerous GRAFS family, also exhibit a significantly increased number of duplications at the basis of vertebrates, together with the Euteleostomi clade (Figure S1A). However, unlike glutamate receptors, their duplication ancestries are of chordate and vertebrate origins (Figure S1C), and they date mostly from the time of WGD at the base of bony vertebrates (Figure S1B).

GPCRs of the rhodopsin class are incredibly diverse and bind a variety of ligand types, including peptides, neuropeptides, hormones, lipids, and ions. Our study indicates that the expansion and diversification of rhodopsin-like GPCRs was not only due to receptor duplication, but largely due to the duplication of their ligands (Figure S1A). These are characterized by a roughly equal number of duplicates and singletons (Table 1) and have duplicated significantly between WGD and the most recent clades up until human speciation (Figure S1A). These duplications arose mainly between Amniota and Theria and were predominantly marked by the expansion of the chemokine signalling system (Figure S1C). Ligands of this class of GPCRs show the youngest duplicate ligands, with most recent duplications arising in anthropoid primates (Simiiformes, Catarrhini and Hominoidea) and belonging to the chemokine, hormone, and neuropeptide categories (Figure S1B). Chemokines are known for their receptor promiscuity, for which gene duplications have made an important contribution (DeVries et al., 2006). This ability to interact with multiple receptors has important biological significance as it increases functional redundancy and promotes flexibility in immune responses, while also allowing for more precise regulation of immunity (Hughes and Nibbs et al., 2018). Another group of ligands of the rhodopsin-like class of GPCRs with an important function in innate immunity are the complement component GPCRs (Yanamadala and Friedlander, 2010). These ligands have an ancient ancestry, and in the present study they were found to date from Euteleostomi and Amniota (Figure S1B). Interestingly, 7TM orphan GPCRs do not share the ancient origin of the five GRAFS families, however, WGD seems to have been particularly important for the expansion of this group (Figure S1A).

G proteins are essential for connecting GPCRs to downstream signalling pathways (Polouliakh et al., 2009; Temple et al., 2010). Given their role as the conserved core of the hourglass network structure, it is unsurprising that they have not undergone recent duplications (Figure 2). This stability reflects their fundamental position in ensuring proper signal transduction across diverse pathways. Our study shows that G proteins are predominantly duplicate genes (Table 1) with ancient origins (Figure S1C), with a very low duplication rate after WGD (Figure S1A), and with most recent duplications dating between WGD and Eutheria (Figure S1B). Their small number and lack of recent duplications is likely related to dosage balance (Birchler and Veitia 2012); in general, signalling systems are dosage sensitive because they are involved in multicomponent interactions, and these have a stoichiometric balance, which may have constrained the evolution of the G *alpha* subunit. Nevertheless, the formation of multiple signalling combinations, such as those involving the different isoforms of the G *alpha* subunit (Figure 3) is considered advantageous, by providing increased diversity and specificity, without changing the modularity of the system (Hur and Kim, 2002). Upon activation by a GPCR, the heterotrimeric G protein dissociates into the G *alpha* subunit and the G *beta gamma* dimer, and these separate parts can act independently to regulate various downstream signalling pathways. Notoriously, the beta and gamma subunits show more recent duplications that were retained until much more recent times in evolutionary history (Figure S1B), suggesting that these two subunits are not so dosage constrained as the subunit *alpha*.

Differences between tissue specificity and duplication frequency depend on interaction types responsible for GPCR activation, and gene expression patterns of the signalling components differ with regard to their evolutionary origins, suggesting that the expansion and evolution of signal transduction in this system involved functional innovations in distinct body systems (Figure 5). Analysis of GPCR subsets with selective ligand binding and G protein coupling profiles revealed that specific interaction types relate to tissue specificity more than to duplication frequency. We also found that the evolutionary origin of gene expression in most tissues was influenced by the duplication history of the signalling components, highlighting the importance of duplication and divergence in the evolution of GPCR signalling in specific tissues and cell types. The analysis of gene expression considering the LDN indicates relevant eutherian origins for most human tissues and important recent evolutionary origins for embryonic structures, which fits the finding that hormones that activate GPCRs are important in human pregnancy (Maston and Ruvolo, 2002), as well as genes involved in pathway crosstalk, which possibly reflect the interaction of GPCRs with other development pathways.

The asymmetric duplication of any interacting component is likely to be detrimental and normally selected against (Gout and Lynch, 2015). Given the early origins of the G protein intracellular transduction system, the duplications that expanded the system are predominantly seen at the receptor level.

In conclusion, this study details the duplication history of the main components of the GPCR signalling system and reveals the importance of the different types of duplication at expanding and diversifying this signalling pathway through its different components at specific evolutionary stages.

## Materials and Methods

### Components of the GPCR signalling system

Human GPCR genes were obtained from IUPHAR/Guide to PHARMACOLOGY (Southan et al., 2015) and filtered to remove all pseudogenes and olfactory receptors. The resulting list was grouped into the <u>G</u>lutamate, <u>R</u>hodopsin, <u>A</u>dhesion, <u>F</u>rizzled/Taste2, and <u>S</u>ecretin (GRAFS) classification system of G protein-coupled receptors (Fredriksson et al. 2003). Ligand interaction information and transduction mechanisms for the GPCR data set were extracted from the same database. Ligands were filtered for the following specifications; species (human proteins), ligand action (agonist only, with the exception of ASIP), and selectivity (natural or endogenous ligands for each GPCR target), and were further classified based on their structure, physiological activity, and GRAFS family they bind to.

Transduction mechanisms were filtered for primary transduction mechanisms only, and the corresponding G proteins were subdivided into their respective subunit classification. A final collection of 1,288 gene-encoded proteins involved in the GPCR signalling system was obtained (Table 1).

### Gene duplication history

Human protein-coding genes were programatically collected from Ensembl version 84 (Zerbino et al., 2017) using the Application Programming Interfaces (API). Ensembl’s Comparative Genomics (Compara) resources were used to classify all human protein-coding genes into duplicates or singletons, according to their evolutionary history. Briefly, Compara’s prediction of gene duplications and speciations is based on phylogenetic trees inferred from 66 vertebrate genomes representing the evolutionary history of gene families that descend from a common ancestor. The gene tree calculation method uses multi-species CDS backtranslated protein-based multiple sequence alignments including the non-vertebrate model species *Caenorhabditis elegans*, *Drosophila melanogaster* and *Saccharomyces cerevisiae*.

Genes for which there are same-species paralogues in the Compara database (i.e., genes of the same species that are related by a duplication event) were defined as duplicate genes. The following were considered singleton genes: 1) genes without gene trees; 2) genes lacking duplication events in their gene trees; or 3) genes without same-species paralogues. Phylogenetic datings for duplicate (singleton) genes were obtained by traversing the gene trees towards the root and recording all taxonomic clades that correspond to duplication (speciation) nodes in the tree (Vilella et al. 2009). The number of duplications in each taxonomic clade was used to estimate duplication ratios across evolutionary time and these were compared to the corresponding proportions for all protein-coding genes as a reference, using a chi-square proportion test. The average number of duplication events for each component of the GPCR signalling system was calculated by dividing all duplication events by the number of genes where the duplication events were found. Significant *P*-values (< 0.05) were adjusted for multiple testing using the Bonferroni method.

### Tissue specificity

RNA-seq data for 27 human tissues were downloaded from the ArrayExpress Archive of Functional Genomics Data (accession number E-MTAB-1733). For all protein-coding genes, tissue specificity was measured by the index τ (Yanai et al. 2004) and by following the protocol described in Kryuchkova-Mostacci and Robinson-Rechavi (2017). A cut-off level of 1 FPKM (fragments per kilobase of exon model per million mapped reads) was set prior to log2 transformation. The FPKM values were averaged for all biological samples, and these values were used to estimate gene expression levels. The τ was calculated using only the genes that were expressed above the threshold in all tissues and according to the following equation;

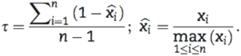

where *n* is the number of tissues, *xi* is the expression of the gene in tissue *i*, and *max(xi)* is the maximum expression level of the gene across all *n* tissues. Gene expression counts were calculated by counting all the tissues where a gene is expressed, using the same threshold chosen for τ.

## Evolutionary analysis of tissue expression

In order to analyse functional organisation within the human body, tissue types were mapped to the Medical Subject Headings (MeSH; http://www.nlm.nih.gov/mesh/) controlled vocabulary under the broad heading Anatomy [A] and classified into their respective systems and structures. Ten pooled organ systems were obtained, for which an expression value was calculated as the average expression within each system.

## Acknowledgements

This work was supported by Fundação para a Ciência e a Tecnologia (SFRH/BD/84834/2012 to A.B.).

## Supplementary Materials

**Figure S1.**
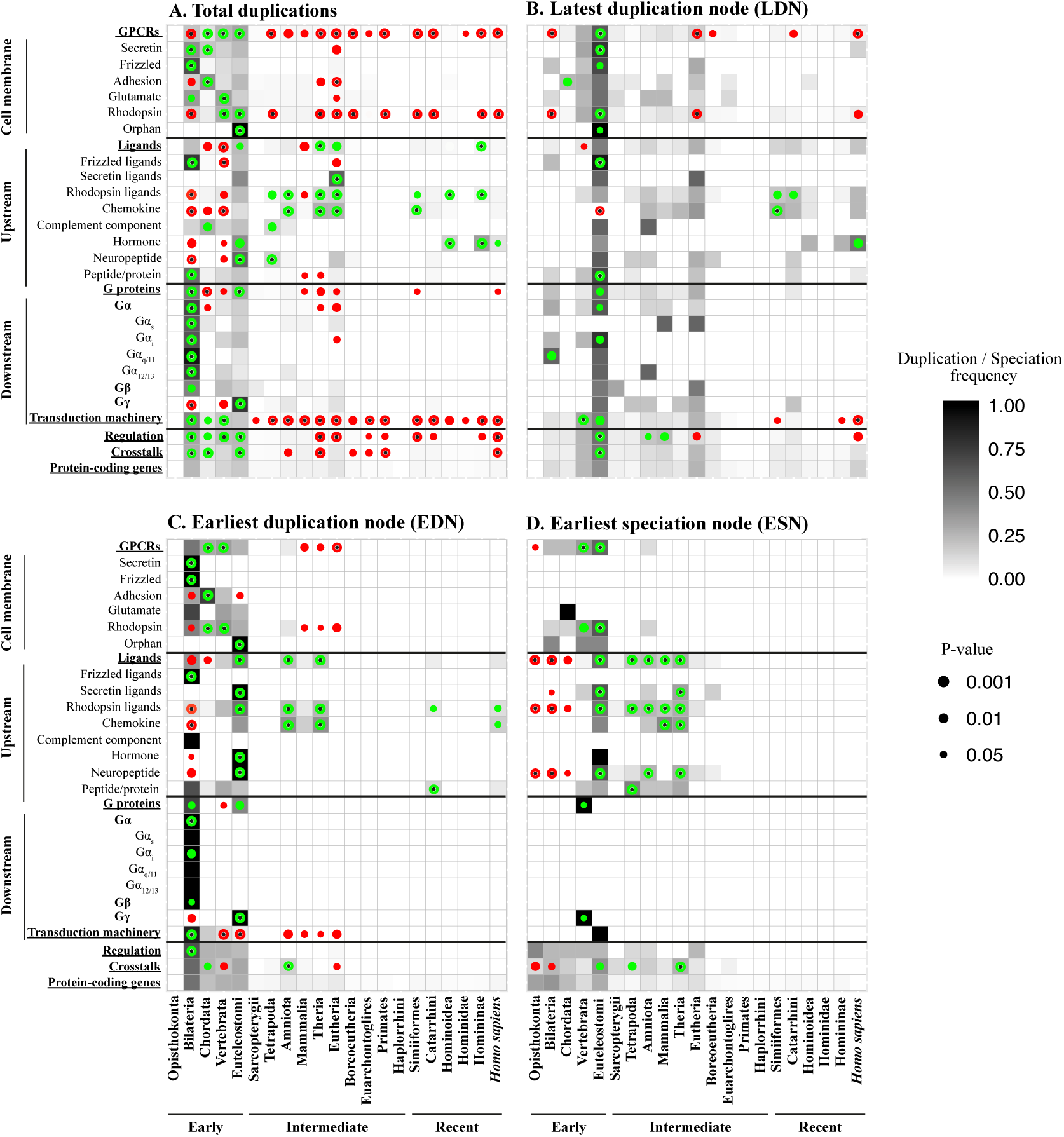
Duplication history of the components of the GPCR signalling system. The grey scale heatmap represents the magnitude of duplication (duplicate genes) and speciation (singleton genes) relative to all protein-coding genes. **(A)** All duplications. **(B)** Latest duplication node (LDN). **(C)** Earliest duplication node (EDN). **(D)** Earliest speciation node (ESN). Circles inside the squares represent significant deviations in relation to all protein-coding genes (*P* < 0.05, χ^2^ test) and green and red colours represent positive or negative deviations, respectively. Black dots inside the circles represent significant *P*-values after Bonferroni correction.

**Figure S2.**
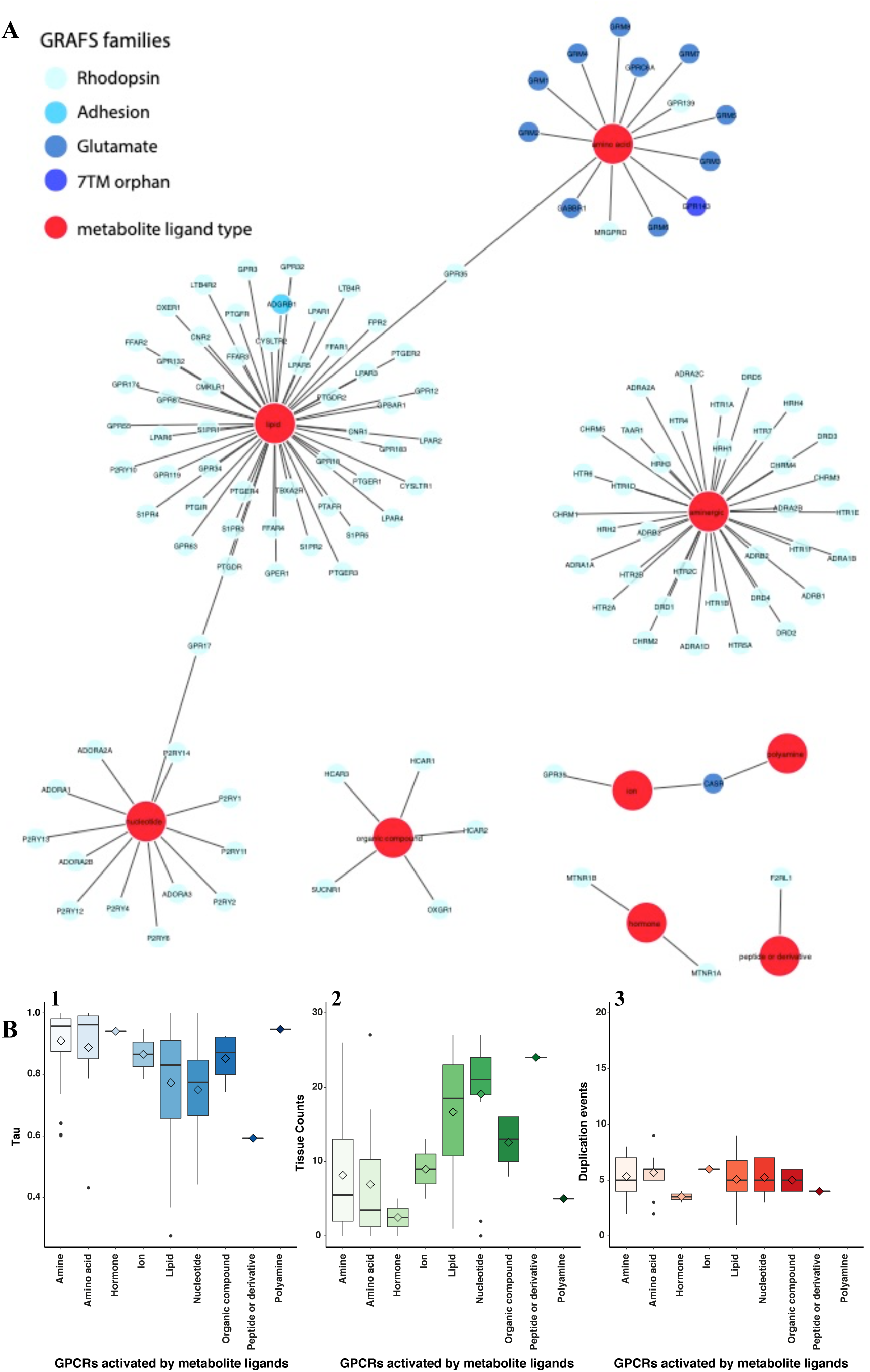
Tissue specificity and duplication events for metabolite-sensing GPCR specificities. **(A)** Edges represent binding interactions between metabolite ligands and their cognate GPCRs. GPCR nodes are coloured according to GRAFS families and the different metabolite ligands are coloured red. **(B).** Boxplots show tissue specificity: (**1)** τ; (**2)** tissue counts; and (**3**) duplication events for each metabolite-sensing GPCR group. Solid lines inside the boxes indicate the median and diamonds indicate the mean.

**Figure S3.**
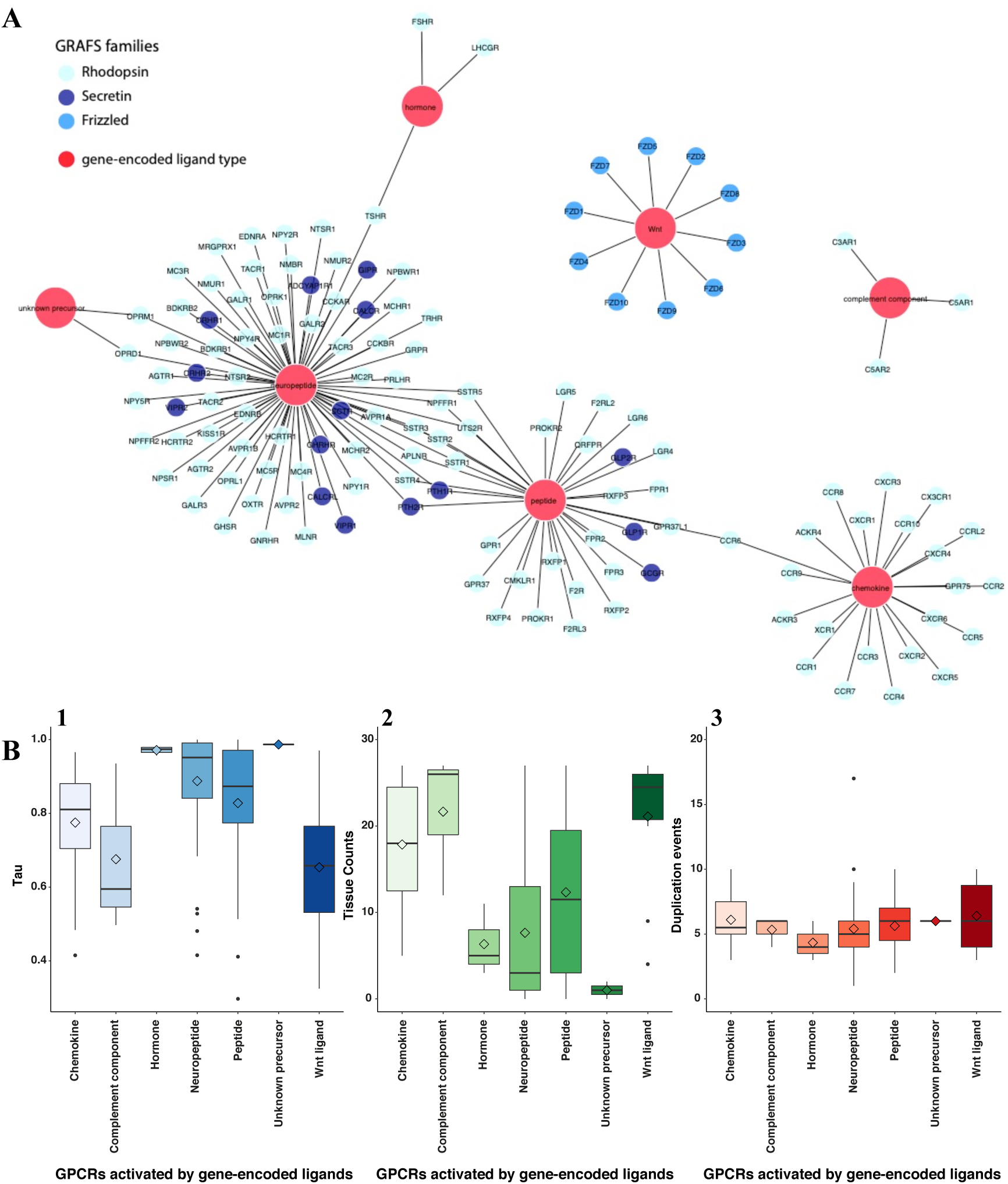
Tissue specificity and duplication events for gene-encoded ligand and GPCR specificities. **(A)** Edges represent gene-encoded ligands and their cognate GPCR binding interactions. GPCR nodes are coloured according to GRAFS families and the different gene-encoded ligands are coloured red. **(B).** Boxplots show tissue specificity: (**1)** τ; (**2)** tissue counts; and (**3**) duplication events for each metabolite-sensing GPCR group. Solid lines inside the boxes indicate the median and diamonds indicate the mean.

**Figure S4.**
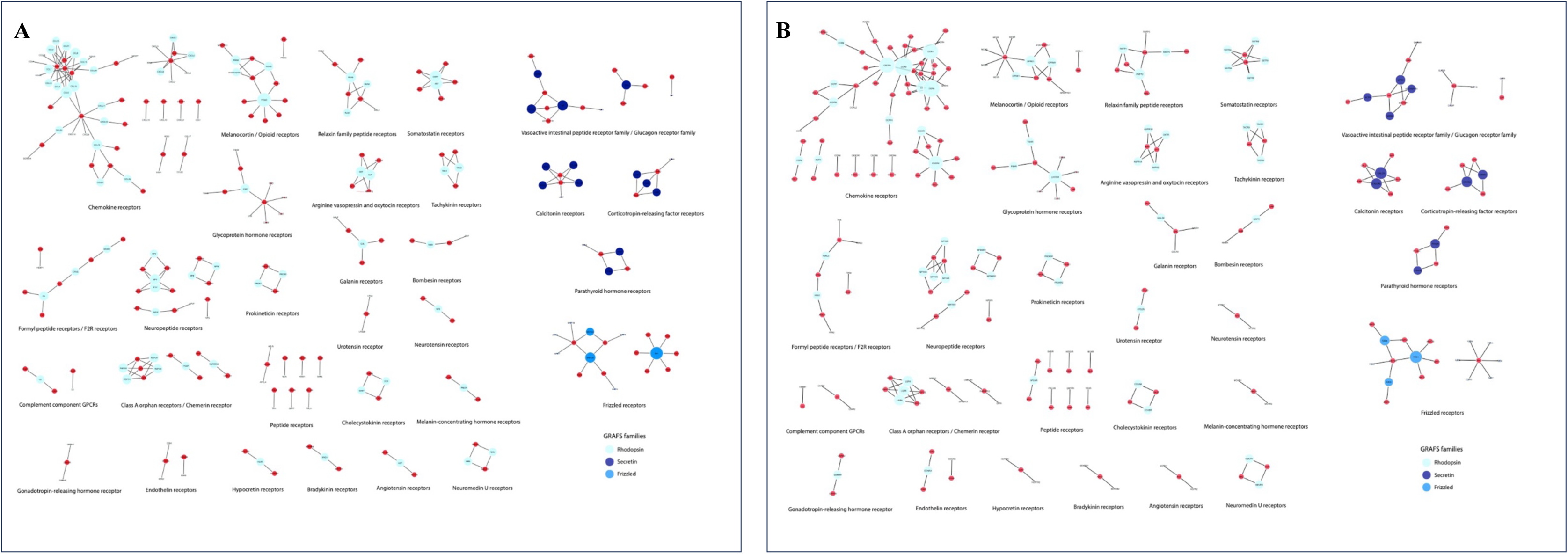
Gene-encoded ligands and their cognate GPCR binding interactions. Edges represent endogenous ligand-receptor pairs. GPCR nodes are coloured according GRAFS families and gene-encoded ligands are coloured red. **(A)** GPCR node size is proportional to the number of agonist ligands activating the receptor. **(B)** Gene-encoded ligand node size is proportional to the number of receptors they activate.

